# PINN-ing the Balloon: A Physics-Informed Neural Network Modelling the Nonlinear Haemodynamic Response Function in fMRI

**DOI:** 10.64898/2026.04.04.716499

**Authors:** Rodrigo H. Avaria, David Ortiz, Javier Palma-Espinosa, Astrid Cancino, Pablo Cox, Rodrigo Salas, Steren Chabert

## Abstract

Accurate characterisation of the haemodynamic response function (HRF) is central to interpreting blood-oxygen-level-dependent (BOLD) signals in functional magnetic resonance imaging, yet standard estimation approaches remain centred around phenomenological formulations lacking biophysical grounding. We present a proof-of-concept methodological study: a physics-informed neural network (PINN) framework that bridges these paradigms by embedding the Balloon– Windkessel model directly into the training objective of a multiheaded neural network. Our approach simultaneously estimates probable latent neurovascular state variables such as cerebral blood inflow, metabolic rate of oxygen consumption, blood volume, and deoxyhaemoglobin content, through an indirect optimisation scheme in which the predicted BOLD signal is obtained via convolution of the estimated HRF with experimental stimuli. Training is governed by a composite loss, balancing differential-equation residuals, hard physics regularisation term, physiological initial conditions and data fidelity. In simulations with temporal signal-to-noise ratios representative of clinical acquisitions, the framework recovered ground-truth state variables with coefficients of determination exceeding 0.95 and mean squared errors below 10^*−*3^, at a physics-to-data weighting of 0.40:0.60. Application to 1.5 T block-design clinical data from an ischaemic stroke patient provides feasibility testing, yielding physiologically plausible, subject-specific HRF estimates, establishing feasibility of single-subject, physics-constrained HRF inference without reliance on fixed gamma basis assumptions. To our knowledge, this constitutes the first deployment of a single PINN incorporating the full Balloon-Windkessel model within an indirect training objective, reconstructing full BOLD observations, thereby positioning PINN-based haemodynamic modelling as a principled and personalised route towards more interpretable and patient-specific biomarkers.

## Introduction

Magnetic Resonance Imaging (MRI) is a cornerstone of clinical and research neuroimaging, as it captures both structural and physiological data, besides being non-invasive. Functional MRI (fMRI) enables the study of brain function *in vivo* by measuring fluctuations in the blood oxygenation level-dependent (BOLD) signal, which arise from haemodynamic and metabolic responses to neuronal activity whether task or state related (1, 2).

The haemodynamic response function (HRF; Fig. 1) is an idealised, noise-free representation of the BOLD response to a brief stimulus (typically *<* 4 s) (3–5). Commonly, the HRF is thought to reflect the brain physiological response, a macroscopic signature of neurovascular coupling shaped by vascular and metabolic dynamics (6). A canonical HRF model arises from studies of haemodynamic responses in healthy individuals and is typically estimated as a combination of two gamma probability density functions, where the first models the shape of the initial stimulus-response, whereas the second models the undershoot (5, 7).

**Fig. 1.**
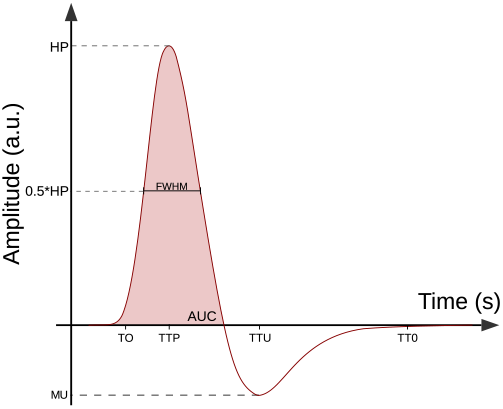
HRF descriptors: (1) Peak Amplitude (Height HP), (2) Time to Peak (TTP), (3) Full Width at Half Maximum (FWHM), (4) Time to Onset (TO), (5) and area under the curve of the first peak (AUC). If there is an undershoot: (6) Minimum Undershoot Height (MU), (7) Time to Undershoot Minimum (TTU), (8) and time to recover baseline value (TT0).

HRF models are crucial for identifying brain activation, and it is often desirable to estimate their parameters with physiological interpretations (8), particularly when exploring the relationship between experimental stimuli and brain responses (9). However, the neural mechanisms linked to metabolism and blood vessel dynamics that generate the BOLD signal remain an active area of research (10, 11).

Various methods and techniques have been proposed to estimate the HRF e.g. Poisson-based approaches (12), Gaussian parameterisations (13), and combinations of gamma functions (14); Bayesian non-parametric estimation (15) and BOLD deconvolution methods that exploit stimulus timing (8, 16, 17).

Several toolboxes support BOLD simulation, e.g., The Virtual Brain, Dynamic Causal Modelling (DCM) in SPM, and NeuRosim (11). In brief, The Virtual Brain typically generates BOLD-like signals by convolving a neural mass/meanfield activity time course with a canonical HRF, whereas DCM uses variants of the Balloon model, a widely used generative model of cerebral haemodynamics (18–21).

The Balloon model, initially proposed by Buxton *et al*. (18) (see Section), represents the venous compartment of a region of interest (ROI) as an expandable “balloon”. The resulting nonlinear dynamical system is commonly written in terms of four normalised state variables that capture key physiological processes: cerebral blood inflow *f* (*t*), cerebral metabolic rate of oxygen consumption *m*(*t*), cerebral blood volume *v*(*t*), and deoxyhaemoglobin content *q*(*t*) (8, 12). A primary challenge, however, lies in precisely identifying this dynamic system, primarily due to noise in experimental data (8).

In recent years, a new type of artificial neural network was developed that leverages purely data-driven methods by incorporating physical equations that describe the process under study. These physics-informed neural networks (PINNs) integrate observational data with mechanistic knowledge by embedding governing equations directly into the training objective (22, 23). The embedded equations constrain learning to physics-admissible solutions, support the solution of forward and inverse problems, and can improve robustness when data are sparse or noisy (24). By leveraging automatic differentiation, PINNs compute derivatives without finite-difference discretisation, which can be advantageous for parameter inference in dynamical models.

From a Machine Learning (ML) perspective, incorporating prior knowledge is a powerful strategy for tackling key challenges, including limited training data, improving model generalisation, and ensuring the physical plausibility of results (24). From a numerical methods perspective, solving differential equations using artificial neural networks (ANNs) differs significantly from classical numerical methods. The solution to the equation comes from optimising the network parameters, which do not depend on the equation’s dimensions, thereby mitigating the problems and drawbacks associated with dimensionality (25).

We present in this work, as a proof of concept, a new PINN-based approach for HRF estimation by combining BOLD observations with the haemodynamic constraints of the Balloon-Windkessel model (12, 18, 20). The method leverages the flexibility of neural networks while enforcing biophysical structure on blood flow, volume, and oxygenation dynamics. In this way, it aims to bridge purely data-driven HRF fitting and fully specified generative haemodynamic modelling, yielding BOLD reconstructions that remain physiologically interpretable.

## Materials and Methods

### The Balloon-Windkessel Model

The Balloon-Windkessel model expresses the BOLD signal *h*(*t*) Eq. (1) as a static, nonlinear function of blood volume *v*(*t*) and deoxy-haemoglobin content *q*(*t*).

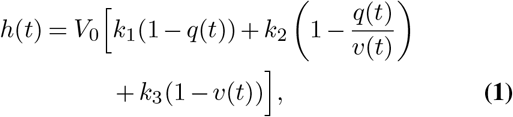

with

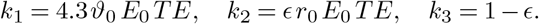

Here *V*_0_ is the resting blood volume fraction; *ϑ*_0_ is the frequency offset at the outer surface of a magnetised vessel for fully deoxygenated blood at 1.5 T; *E*_0_ is the resting oxygen extraction fraction; *TE* is the echo time; *ϵ* is the ratio of intra- and extravascular signals; and *r*_0_ is the slope relating the intravascular relaxation rate 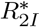 to oxygen saturation. Although alternative parameter values and model extensions can be found in (19–21, 26), we adopt the values of *k*_1_, *k*_2_, and *k*_3_ reviewed by Stephan *et al*. (21), which have been used in subsequent work (11, 27–32).

As shown in Eq. (2), and given a mean transit time *τ*_*MTT*_, the dynamics of *v*(*t*) are driven by inflow *f* (*t*) through an outflow term *f*_*out*_(*v, t*). The deoxyhaemoglobin state *q*(*t*) is jointly shaped by blood volume, outflow, and the metabolic drive *m*(*t*) (18).

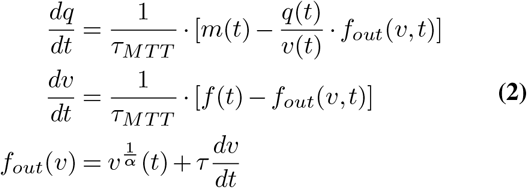

Here *α* is the Grubb stiffness exponent (a proxy for venous compliance) (21, 33), and *τ* controls the time scale of the viscoelastic outflow adjustment.

We follow the two-input pipeline in Fig. 2 proposed by Maith *et al*. (11), which builds on extensions suggested by (12) and (10). The key modelling choice is to separate the inflow *f* (*t*) from the metabolic drive *m*(*t*), allowing them to be driven in parallel by the same stimulus input *I*(*t*) through Eq. (3).

**Fig. 2.**
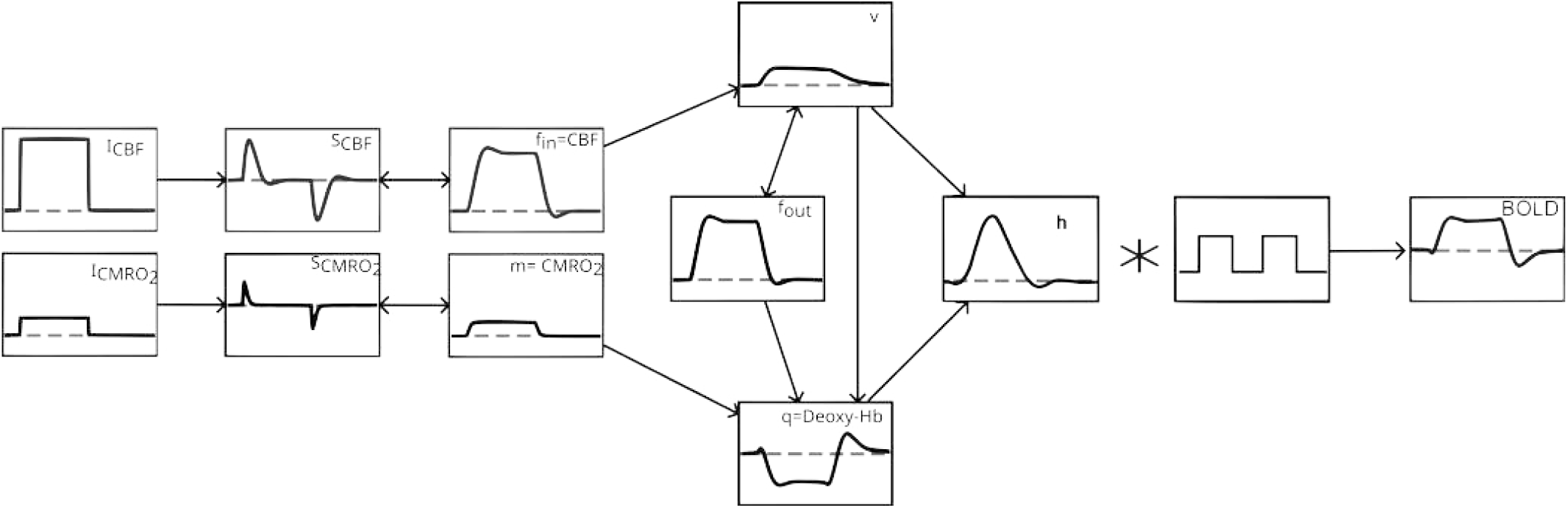
Balloon model with parallel-driven *CBF* and *CMRO*_2_, coupling the increase in q with the normalised *CMRO*_2_ (m) directly (see (20)), which is driven by a second input signal 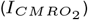 like the normalised *CBF* (increased to 0.05). Further, *f*_*out*_ is described by the equations of Buxton *et al*. (20), which causes v to decrease more slowly.

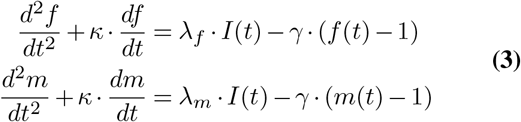

We choose *κ* and *γ* such as *f* (*t*) and *m*(*t*) exhibit an under-damped response (allowing both overshoot and undershoot), and *λ*_*f*_ *>> λ*_*m*_ making m’s oscillation smaller in amplitude.

### Physics-Informed Neural Network

A standard PINN architecture comprises an input layer, one or more hidden layers, and an output layer. The input is a feature vector sampled from the problem domain; the hidden layers learn a latent representation; and the output layer produces the final prediction vector (34, 35).

Within this framework, we represent the unknown solutions of Eq. (2) and Eq. (3) with a single deep neural network **u**_*θ*_(*t*), where *θ* denotes all trainable parameters (weights and biases). We implement a multi-headed (parallel-output) multilayer perceptron, adapted from Protopapas and Raval (34), to simultaneously produce the four Balloon state variables. Our model (Fig. 3) takes a normalised, uniformly sampled time vector 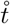 as input (mean-centred, unitary variance 30 s, sampled every centisecond), uses two hidden layers (128 *→* 256 units, 256 *→* 512 units), and applies Softmax activations in the shared stem. The final hidden layer is split into two equally sized 256-neuron buds, before connecting to the output layer. Although each network instantiation has random initial *θ* values according to Glorot initialisation (36), we implement factorised the layers in the shered stem according to Wang *et al*.(37, 38) random weight factorisation *i*.*e*. for every layer *l* = 1, 2, 3, and its initial weight matrix *W* ^*l*^ we proceed by initialising a scale vector *exp*(*s*) where *s* is sampled from a multivariate normal distribution *N* (*µ, σI*) and rewrote *W* ^*l*^ = *diag*(*s*^*l*^) *· V* ^*l*^. During training gradient descent is applied directly to the parameters *s* and *V*. The output layer comprises four linear “heads” (Softplus activation with *β* = 1.0 and threshold = 20.0), corresponding to the Balloon state variables. One of the 256-neuron buds connects to **u**_**f**_ and **u**_**m**_ (256 *→* 1 each), and estimate *f* (*t*) and *m*(*t*) (as 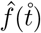 and 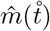 respectively) from Eq. (3). The connections amog the heads mirrors the balloon model system of equations, thus **u**_**f**_ joins with the second 256-neuron bud when connecting with **u**_**v**_ (257 *→* 1), who estimates *v*(*t*) from Eq. (2) 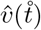. **u**_**v**_ and **u**_**m**_ join with the second 256-neuron bud tribute into **u**_**q**_ (258 *→* 1) estimating *q*(*t*) 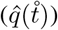 from Eq. (2).The network output is a 4 *×* 3000 array.

**Fig. 3.**
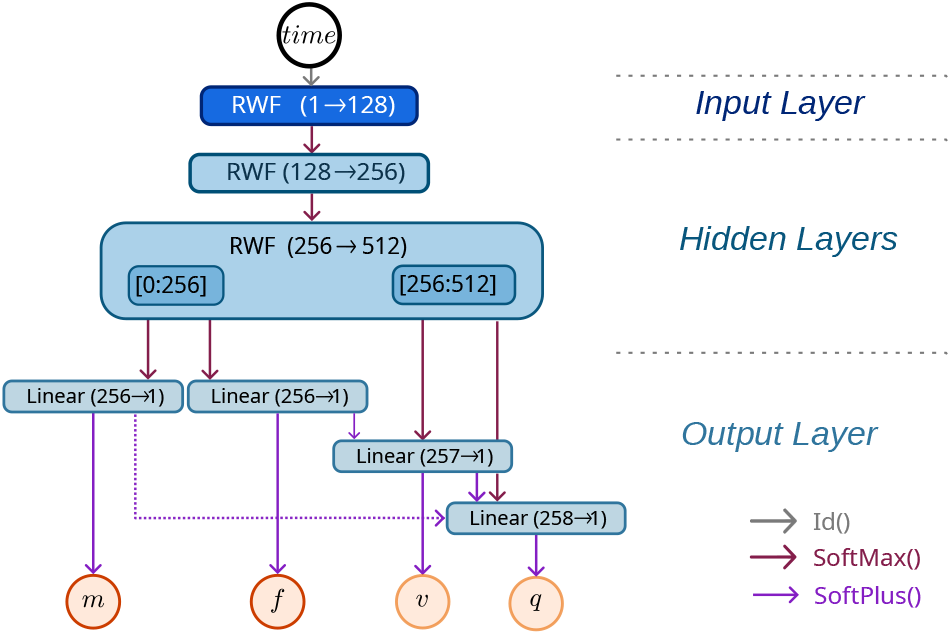
PINN Architecture: A normalised time coordinate enters a Random Weight-factorisation (RWF) input layer. We used a two RWF hidden-layer network architecture. The RWF layers processes its inputs through a Softmax activation function (dark red arrows). The output of the hidden layer is split into two equally sized 256-neuron buds([0:256] and [256:512]), before feeding into the output layer. The output layer has 4 linear heads with Softplus activation function (purple arrows), mapping to the state variables: **û**_**m**_ and **û**_**f**_ read the first bud obtaining 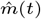and 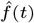 reads the second bud in addition to 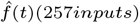, whereas **û**_**q**_ reads also the second bud jointly with 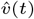 and 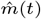 (thus the 258 inputs). Blue boxes indicates the layers; Orange circles indicate the output, estimated state variables.

Softmax election was motivated by its low dimension resembling the hyperbolic tangent (Tanh), selected among smooth (*C*^*∞*^) candidates given the second-order ODE residual to minimise and for its bounded, normalised output.

The HRF is not a direct network output. Instead, it is obtained by substituting 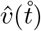 and 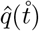 into Eq. (1), the resulting 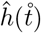 is then used to reconstruct the BOLD signal via Eq. (5). We will use 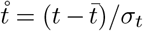 for the zero-mean, unit-variance rescaling of physical time onto [ *−*1.73, 1.73], chosen to improve gradient conditioning.

A central appeal of PINNs is that they are mesh-free: the solution is represented by a neural network, and training reduces to an optimisation problem, i.e., finding the network parameters 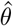 that minimise a total loss *L*_*tot*_ Eq. (4). This avoids the grid whose size grows exponentially with the dimension of the problem (Raissi *et al*. (24)).

When PINNs are used to solve differential equations, *L*_*tot*_ is typically defined as a weighted sum of a data-fitting term *L*_*data*_, an equation-residual term *L*_*eq*_, and an initialcondition term *L*_*ic*_. In our experiments, added measurement noise degraded a fraction of the fits; to stabilise training and reduce these implausible solutions, we introduce a fourth term, a physics-regularisation term *L*_*reg*_, giving Eq. (4)

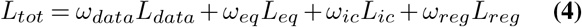

We adopt an indirect objective for the data-fitting term *L*_*data*_. The predicted BOLD signal *Ŷ* (*t*) is obtained by convolving the estimated HRF *ĥ* with the stimulus function *s*(*t*) Eq. (5); the prediction is subsampled to the acquisition repetition time (TR) and compared with the observed samples 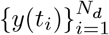 (simulated or *in vivo*) through the mean squared error Eq. (6).

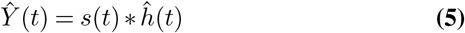

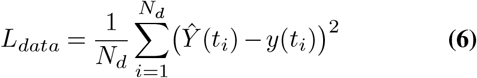

The equation-residual term *L*_*eq*_ enforces the Balloon-model dynamics: the unknown solution is represented by the network, which lets us define the ordinary differential equation (ODE) residuals as

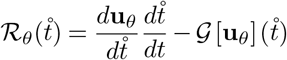

where *G* is a differential operator; evaluating the residuals in this standardised domain requires reparameterising the Balloon ODEs according to the chain rule (full derivation in Appendix 2). Thus, minimising these residuals yields the physics loss *L*_*eq*_, parametrised by 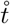. Then *L*_*eq*_ can be expressed as

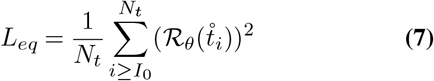

In our case, the Balloon model is formulated as a system of ordinary differential equations; thus, 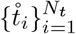 denotes the set of standardised time-domain samples. To solve Eq. (3), we apply a single brief impulse (a 1 s boxcar). Its onset index *I*_0_ marks the end of the resting baseline period used to impose initial conditions. Concretely, we constrain not only the states at *t* = 0 (i.e., *f* (0), *m*(0), *v*(0), *q*(0)) but also all samples with 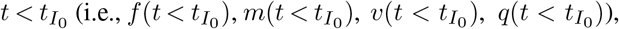, which encourages the network to maintain physiologically plausible basal levels before stimulation. When solved numerically, Eq. (3) is typically reduced to an equivalent first-order system and integrated with Dirichlet (state-only) initial conditions. Because the PINN retains the second-order formulation directly, Cauchy conditions are the initial conditions imposed here, constraining both the state and its first derivative for all 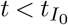.

Equation Eq. (8) encodes Cauchy’s initial condition: the first term penalises the network output deviations from the baseline state, and the second term penalises deviations of its first derivative from the baseline derivative, both evaluated for 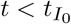; together, they uniquely specify the solution of the second-order system. The quantities *x*_0_ and **x**_0_ denote the baseline values for the state variables and their derivatives, respectively.

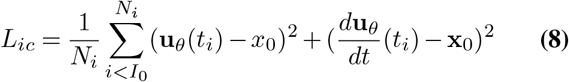

The physical regularisation term comes from Eq.9, an outflow-free, exact dynamic constraint on the Balloon state pair.

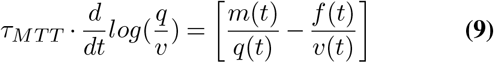

The expression makes no assumption about a specific venous compliance of the model. Without committing to the outflow submodel, it constrains a combination of states/derivatives in a single expression; its full deduction can be seen in Appendix 3.

Making use of Eq. (9) we state 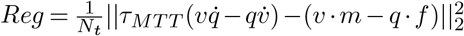, as a hard regularisation term.

Therefore, the PINN must solve the dynamical system in Eq. (2), Eq. (3) and Eq. (9) while simultaneously explaining the observed BOLD signal through the forward observation model in Eq. (1) and Eq. (5).

### Experiments

Our aim is to assess PINN’s ability to estimate the HRF by regressing the convolution of the balloon model’s output with known stimuli against an observed BOLD signal. This requires solving the coupled ODE systems in Eq. (2) and Eq. (3), before the convolution.

We evaluate the performance under three experimental settings. First, retrieving the HRF from noiseless BOLD Simulated Data. Secondly, having verified that our proposal is able to restore the Balloon model state variables under ideal conditions, we proceeded to retrieve the HRF from noisy BOLD simulated data: in these experiments, we trained our PINN following the previous paradigm, introducing additive Gaussian noise before subsampling to emulate the target temporal signal-to-noise ratio (tSNR) and TR conditions. Thereafter, we applied our framework to real BOLD data from a previously performed block-designed fMRI study, training the PINN against *in vivo* data.

Given that the clinical experiment lacks physiological ground truth, this last assessment is presented as a feasibility study demonstrating convergence and physiological plausibility under everyday clinical conditions; however, it cannot serve as validation of physiological latent-state recovery.

Due to random *θ* initialisation of the network, each experiment was performed 100 times, yielding 100 HRF estimates *ĥ*(*t*). Each network instantiation has random initial *θ* values according to Glorot initialisation (36), every training was performed with ADAM optimiser with 6050 iterations, an initial learning rate of 1 *×* 10^*−*3^, and a step-wise decay factor of 0.15 every 1000 iterations. The loss function weights proportion *ω*_*data*_*/ω*_*ode*_*/ω*_*ic*_*/ω*_*reg*_ were kept constant at 0.60 : 0.40 : 1.0 : 1.0 for all experiments.

Following (39), we characterise each *ĥ* (*t*) using the descriptors shown in Fig. 1: height to peak (HP), time to peak (TTP), FWHM, time to onset (TO; time to a 10% increase from baseline), and the area under the first peak (AUC; using trapezoidal rule). When an undershoot is present, we also report the minimum undershoot height (MU), time to undershoot minimum (TTU), and time to return to baseline (TT0). After visual inspection, we validated the results by identifying implausible descriptor values, following empirical criteria similar to those defined by (14) and used by (39): any descriptor value outside predefined acceptable ranges will deem its HRF, and state variales, estimation abnormal and therefore be excluded from subsequent analysis. The criteria considered include, but are not limited to, peaks occurring during or before the impulse function I from Eq. (3), nought HP, a BOLD peak signal of magnitude greater than 15%, *TTP >* 15 s, FWHM greater than 10 s, or TO lower than 1.5 s.

All experiments were implemented in Python 3.10.12 using `PyTorch` 2.3.1 and `CUDA` 12.1. Computations were performed in single-precision floating-point representation on a single NVIDIA GeForce RTX 4080 GPU, on 13th Gen Intel Core i9-13900 x 32. Ubuntu 24.04.4 LTS, GNU-Linux 6.8.0106-generic as OS. All codes used to perform these experiment are bundled together in a single python library avaliable at Rodrigo H. Avaria’s github

#### Bayesian equivalence assessment of descriptor recovery

Each descriptor is estimated from an ensemble of *n* = 100 random-initialisation fits; in this condition, a point-null test of “zero bias” rejects for offsets far smaller than the measurement resolution: with *n* this large, the null is almost always rejected, and the p-value reports precision rather than relevance. We therefore estimated the bias of each descriptor and assessed whether it was practically zero using Bayesian estimation with a Region of Practical Equivalence (ROPE) (40). For descriptor *d* with true value 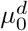 (obtained by applying the descriptor operator to the ground-truth HRF), we modelled the *n* ensemble values as Normal with unknown mean and variance under the non-informative Jeffreys prior *p*(*µ, σ*^2^) *∝ σ*^*−*2^. This yields a closed-form marginal posterior for the bias 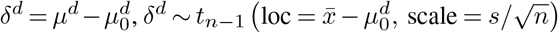, a Student-*t* distribution with *n −* 1 degrees of freedom, where 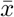 and *s* are the ensemble mean and sample standard deviation. No MCMC or variational approximation is required; the posterior is exact and evaluated analytically.

From this posterior we report the 95% highest-density interval (HDI; equal-tailed and thus the HDI for the symmetric *t*) and compare it to a ROPE of half-width Δ^*d*^ chosen per descriptor to reflect the physiological/temporal resolution of interest: an absolute margin of *±*0.1 *s* for the time descriptors (TTP, FWHM, TO, TTU, TT0) and *±* 1.5% of |*µ*_0_| for the dimensionless/amplitude descriptors (HP, AUC, MU). Each descriptor was assigned a three-way verdict by Kruschke’s HDI-and-ROPE rule: equivalent if the HDI lies entirely inside [*−*Δ, Δ], biased if it lies entirely outside, and undecided if it straddles a ROPE limit.

#### Simulated data

In the first two experiments, we train the PINN using simulated data generated through a custom implementation of the Balloon model, with an integration step *dt* = 0.01. We simulate the response to a single 1 s impulse *I*(*t*) in Eq. (3), propagate the resulting states through Eq. (2), to compute *h*(*t*) using Eq. (1), and finally generate BOLD time series by convolving *h*(*t*) with a 288 s stimulus train, equivalent to an experimental protocol comprising ten 3 s blocks separated by irregular intervals. Parameter values for Eq. (1), Eq. (2), and Eq. (3) follow (11, 20) and are summarised in Table 4. After the balloon simulation and convolution, we subsampled *Y* (*t*) every 175 samples to mimic 165 volumes (*TR* = 1.75 s; approximately 5 min acquisition).

For the second simulation setting, following (14), we add independent Gaussian noise to achieve a prescribed tSNR equivalent to *in-vivo* conditions

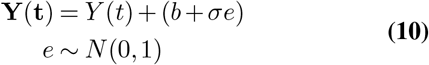

Here **Y**(**t**) corresponds to the noise-added simulated fMRI data; *Y* (*t*) denotes the ideal noiseless BOLD simulation from the convolution, whereas *b* = 116.8 corresponds to the base level and *σ* = 1.45 to the additive noise standard deviation in order to emulate a real *tSNR* ∼ 67. After simulation and noise addition, we subsampled **Y**(**t**) similarly to the first experiment to mimic 165 volumes (*TR* = 1.75 s; *tSNR* = 64; approximately 5 min acquisition).

### Application to *In Vivo* Data

We next apply the model to observational in vivo data to appraise HRF recovery from a signal with real clinical conditions. Since physiological ground truth is unavailable, we do not aim to validate the recovered latent states. This experiment should be interpreted as a feasibility assessment illustrating whether the PINN can converge to physiologically plausible solutions when confronted with real clinical BOLD data.

Data come from a study on acute ischaemic stroke conducted between March 2022 and September 2023, approved by the Regional Ethics Committee under Resolution *N o*.200 *−* 2026, and carried out following the 2013 revision of the Declaration of Helsinki. The patient, a 52-year-old male, presented with an ischaemic core in the right thalamic region measuring 0.55 ml in volume, with a National Institutes of Health Stroke Scale (NIHSS) score of 1 and an evolution time of 7 hours before hospital arrival.

Images were obtained using an eight-element head coil on a 1.5*T* Signa HDxt scanner (General Electric, Milwaukee, WI, USA). For fMRI, *T* 2^*\**^-weighted echo-planar imaging (EPI) was employed with a TR of 1.75 s, echo time (TE) of 60 ms, and a spatial resolution of 1.9 *×* 1.9 *×* 5*mm*^3^. The stimulation paradigm involved passive bilateral wrist flexion-extension movements (41), manually performed by a third party at approximately 1*Hz*. Movement timing was coordinated using visual cues on a screen, signalling the start and stop of each motion. Each wrist movement lasted 3 seconds, with interstimulus intervals ranging from 15 to 30 seconds, yielding 11 event-related activations. The total acquisition time for the fMRI session was 5 minutes.

After standard preprocessing, slice timing correction was applied, followed by realignment (estimation and reslicing), co-registration of functional and anatomical images, and smoothing with a full width at half maximum (FWHM) of 6 mm in each direction (6, 6, 6) using **SPM12** software (Wellcome Trust Centre for Neuroimaging, London, UK). The postcentral gyri were defined as the regions of interest (ROI) for BOLD signal extraction. The ischaemic lesion side of the patient determined the designation of the ischaemic postcentral gyrus (PCG) and the non-ischaemic PCG. We applied baseline correction by subtracting the mean BOLD signal value from the whole time series.

## Results

### Noiseless Simulation

The time series estimated after 6050 training iterations for each state variable are illustrated in Fig. 4, within their standard deviation; no estimations were excluded, thus we compare 100 realisations of each estimation: normalised blood inflow, cerebral metabolic rate of oxygen, volume, and deoxyhaemoglobin concentration time series predicted by the PINN model against the numerical solutions (our “ground truth”), finding reasonable agreement between them when measured using Coefficient of determination (*R*^2^), *L*_2_ relative error (*L*_2*RE*_) and Spearman Correlation (*Sρ*) (Table 1).

For our experiment on BOLD noiseless simulated data, we obtained a mean squared error (MSE) lower than 10^*−*4^. The *L*_2*RE*_ between our estimations and the ground truth signal was ~ 10^*−*3^. The MSE between the reconstruction and the data were ~ 10^*−*5^. Performance metrics, for all hundred ensembles of training runs, are summarised in Table 1. We observe that our network closely approximates the numerical state variables; a comparison of its HRF descriptors (shown in Fig. 1), is shown in Fig. 5 where whole set of descriptors can be seen plotted against a *±*0.1 s ( *±*1.5% for HP/AUC/MU) ROPE in Fig. 5.

**Table 1.**
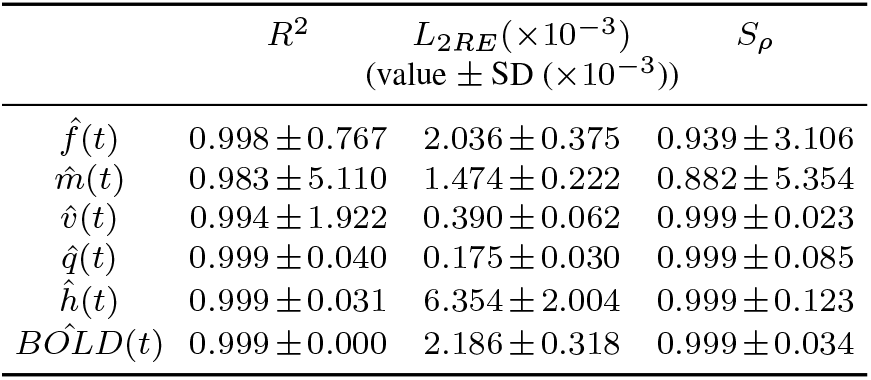
Balloon state variables regression and BOLD reconstruction mean performance metrics against noiseless simulated data, for *n* = 100 ensemble, random-initialisation and their standard deviation (SD). *R*^2^ is the coefficient of determination, *L*_2*RE*_ the relative error, and *S*_*ρ*_ the Spearman correlation coefficient. SD values for *R*^2^ and *L*_2*RE*_ are *×*10^*−*3^; those for *S*_*ρ*_ are *×*10^*−*2^.

| | $R^2$ | $L_{2RE}(\times 10^{-3})$<br>(value ± SD ($\times 10^{-3}$)) | $S_\rho$ |
| --- | --- | --- | --- |
| $\hat{f}(t)$ | $0.998 \pm 0.767$ | $2.036 \pm 0.375$ | $0.939 \pm 3.106$ |
| $\hat{m}(t)$ | $0.983 \pm 5.110$ | $1.474 \pm 0.222$ | $0.882 \pm 5.354$ |
| $\hat{v}(t)$ | $0.994 \pm 1.922$ | $0.390 \pm 0.062$ | $0.999 \pm 0.023$ |
| $\hat{q}(t)$ | $0.999 \pm 0.040$ | $0.175 \pm 0.030$ | $0.999 \pm 0.085$ |
| $\hat{h}(t)$ | $0.999 \pm 0.031$ | $6.354 \pm 2.004$ | $0.999 \pm 0.123$ |
| $BOLD(t)$ | $0.999 \pm 0.000$ | $2.186 \pm 0.318$ | $0.999 \pm 0.034$ |

**Fig. 4.**
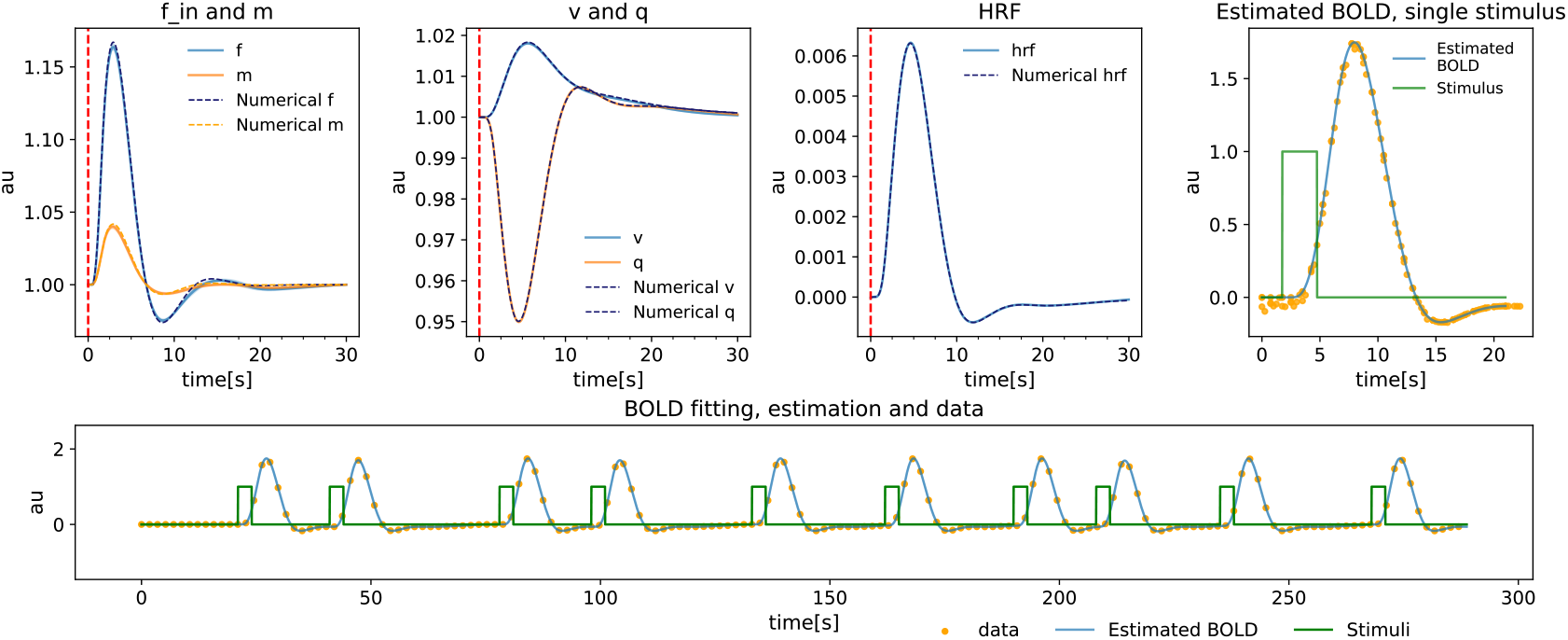
Ensemble reconstruction of the Balloon state variables and the haemodynamic response from synthetic noiseless BOLD data. *n* = 100 independent PINN restarts were trained on the same time series; every panel shows the across-restart mean (solid) with a *±*1 SD band, and the dashed curves give the numerical solution of the forward model used to generate the data. From left to right Inflow *f*_*in*_ and metabolic rate *m* from Eq.3; venous volume *v* and deoxyhaemoglobin content *q* from Eq.2; the recovered impulse response *h*(*t*) from using Eq.1. The vertical dashed line marks stimulus onset. Single-stimulus BOLD prediction obtained in Eq.5 by convolving each restart’s HRF with one stimulus block (green), overlaid on the stimulus-locked epochs of the measured signal (orange). Second row: the ensemble BOLD reconstruction against the complete noise-free recording across the ten-block stimulus train. The recovered HRF peaks at *HP* (6.328 *±* 0.013) *×* 10^*−*3^ a.u. at 4.620 *±* 0.02 s, has a FWHM of 4.776 *±* 0.012 s, and reaches an undershoot MU of (*−*6.26 *±* 0.05) *×* 10^*−*4^ a.u. at 11.77 *±* 0.05 s (mean *±* SD, *n* = 100). All quantities are in arbitrary units; time in seconds.

**Fig. 5.**
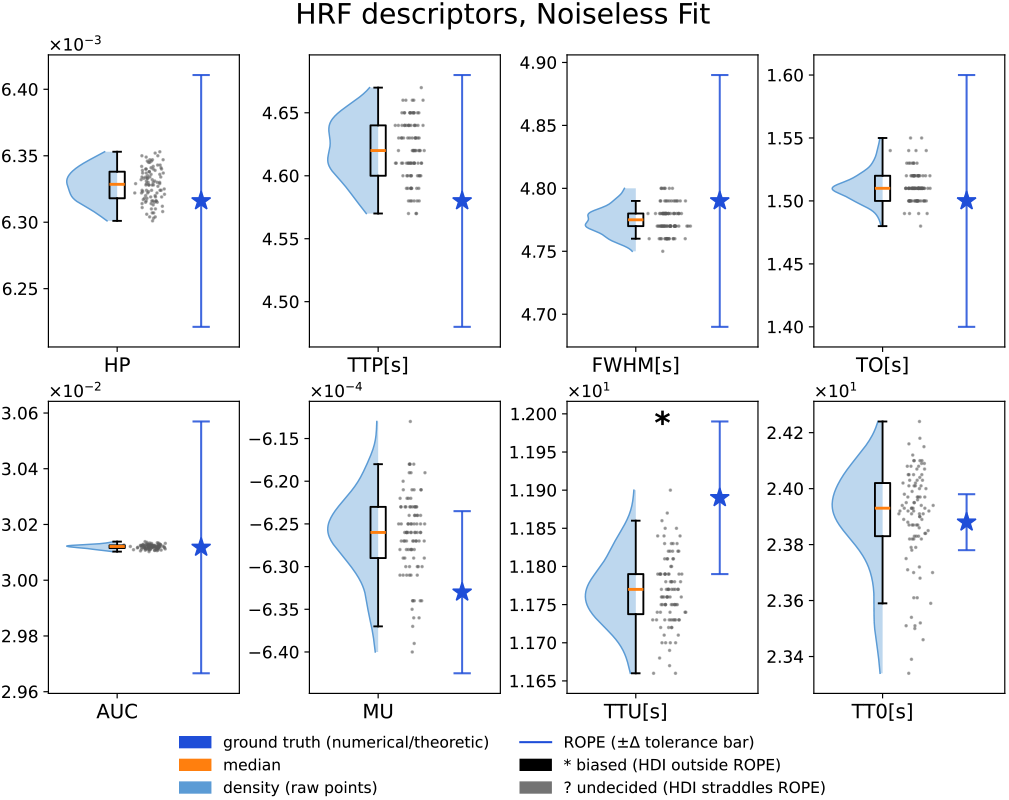
HRF descriptors across the noiseless ensemble (n=100 independent PINN fits of the same noise-free simulated BOLD signal). Each panel shows one descriptor as a raincloud: a half-violin kernel-density estimate of the ensemble (light blue), a box spanning the interquartile range with an orange mark as the median, and jittered gray raw points for the individual runs. The blue star marks the descriptor’s value for the ground truth from the numerical HRF, the vertical blue line spans the ROPE (*±*Δ), with Δ of 0.1 *s* for the time descriptors (TTP, FWHM, TO, TTU, TT0) and *±*1.5% of |*µ*_0_| for the dimensionless/amplitude descriptors (HP, AUC, MU). For each descriptor bias is estimated from an analytic conjugate Student-t posterior under a Jeffreys prior and classified by the HDI-versus-ROPE rule at 95% credible mass. Seven of the eight descriptors are recovered within the stated tolerance; only TTU is flagged as biased; the ensemble has a small, though systematic, offset *≈* 0.12 *s* below truth. Note the per-panel axis multipliers.

Seven of the eight descriptors were practically equivalent to ground truth: their 95% HDIs fell entirely within the equivalence margin. TTU was the sole descriptor judged biased (*−*0.122 s) with its 95% HDI lying wholly below the ROPE limit, with *<* 0.01% of the posterior inside the ROPE. The noiseless ensemble therefore recovers the HRF shape to within *±*0.1 *mathrms* on every descriptor except a small, well-resolved ~ 0.12 s early bias in the undershoot timing (Table 5, Appendix 4).

### Noise-Added Simulation

Having trained the model anew 100 times, each state variable time series estimated by PINN is illustrated in Fig. 6, next to the standard deviation. No training run was excluded. Therefore, as before, we compare 100 state variable estimations against the numerical “ground truth”, this time under additive noise (*tSNR* = 64), finding reasonable agreement (Table 2). We observe that, with a weight ratio of 0.40 : 0.60 between physics and data, our PINN model is capable of closely approximating the numerical state variables. The corresponding performance metrics are shown in Table 2, whereas the estimated state variables are shown in Fig. 6.

**Table 2.** Balloon state variables regression and BOLD reconstruction mean performance metrics (and their standard deviation, SD) against simulated data with white noise added, *n* = 100 randominitialisation, *tSNR* = 64. *R*^2^ the coefficient of determination; *L*_2*RE*_ relative error and *S*_*ρ*_ Spearman Correlation coefficient

| | $R^2$ | $L_{2RE}(\times 10^{-3})$<br>(value ± SD ($\times 10^{-3}$)) | $S_\rho$ |
| --- | --- | --- | --- |
| $\hat{f}(t)$ | $0.999 \pm 0.235$ | $1.130 \pm 0.196$ | $0.911 \pm 3.084$ |
| $\hat{m}(t)$ | $0.990 \pm 2.414$ | $1.182 \pm 0.139$ | $0.890 \pm 1.850$ |
| $\hat{v}(t)$ | $0.957 \pm 1.662$ | $1.089 \pm 0.161$ | $0.995 \pm 0.107$ |
| $\hat{q}(t)$ | $0.991 \pm 2.422$ | $1.588 \pm 0.207$ | $0.962 \pm 1.836$ |
| $\hat{h}(t)$ | $0.994 \pm 1.698$ | $72.444 \pm 9.862$ | $0.958 \pm 45.22$ |
| $BOLD(t)$ | $0.256 \pm 1.204$ | $86.26 \pm 7.011$ | $0.408 \pm 14.29$ |

**Fig. 6.**
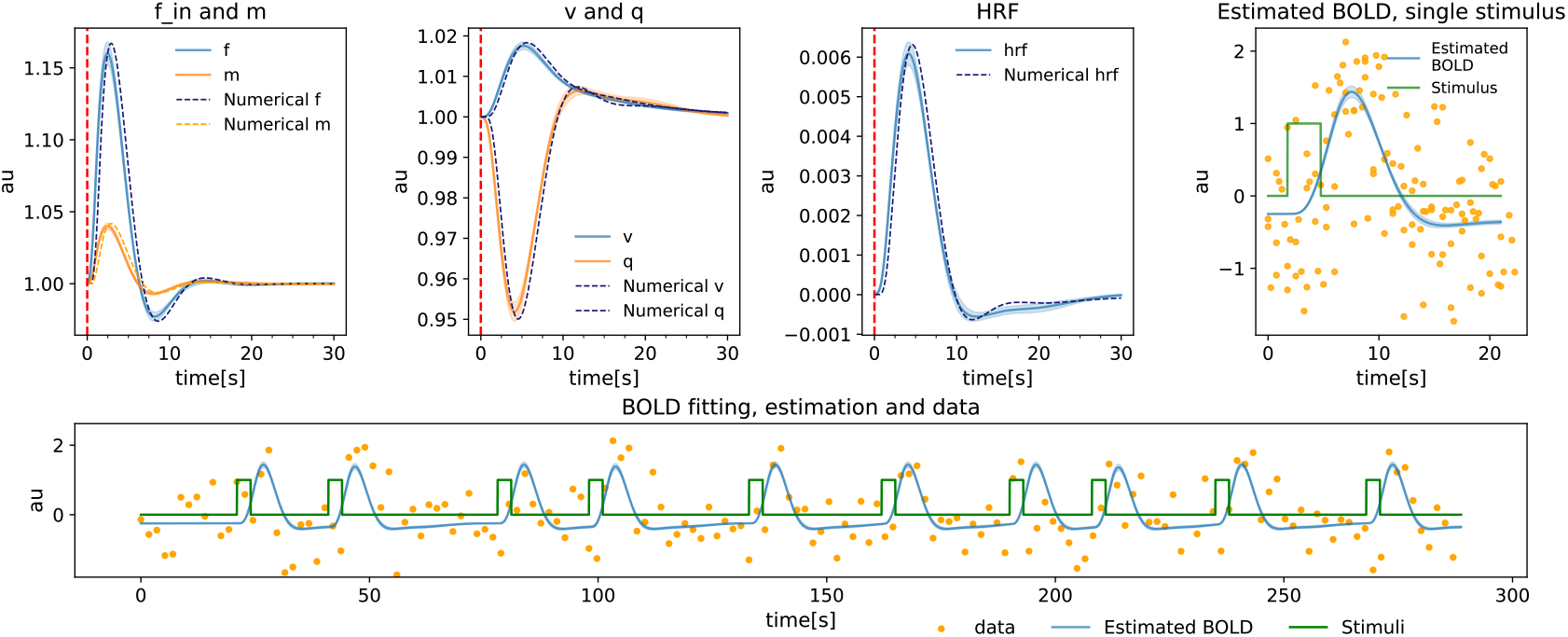
Ensemble reconstruction from noise-added BOLD data (tSNR=64; *n* = 100 PINN restarts). Mean (solid) *±*1 SD across restarts; dashed curves are the numerical forward solution. Left to right, top row: *f*_*in*_ and *m* from Eq.3; *v* and *q* from Eq.2; the recovered HRF using Eq.1, HRF peak (6.272 *±* 0.065) *×* 10^*−*3^ a.u. at *≈* 4.60 s.; and a single-stimulus BOLD prediction using Eq.5 against the stimulus-locked epochs of the scattered measurement (orange). Bottom row: the full-session fit over the ten-block stimulus train. The across-restart SD never exceeds 1.5% of a state’s dynamic range, so the offsets from the reference reflect systematic bias, not initialisation variance; the low BOLD *R*^2^ in Table 2 reflects the unpredictable noise component, not disagreement among restarts.

Fig. 7 shows that under a *±*0.1 s ( *±*1.5% for HP/AUC/MU) ROPE, four of the eight descriptors were practically equivalent to ground truth (datailed in Table 6, (Appendix 4)): their 95% HDIs fell entirely within the equivalence margin (e.g., TTP bias +0.017 s, 95% HDI [+0.012, +0.021]; FWHM bias 0.07 s, 95% HDI [+0.066, +0.075]). AUC was the sole descriptor judged undecided: its bias of +0.00051 (1.7%) had a 95% HDI of [+0.00045, +0.00058], slightly above the 1.5 % ROPE limit, with *<* 2.8% of the posterior inside the ROPE. The ensemble therefore recovers the HRF shape to within *±*0.1 s for the first half of process but falls outside the ROPE for the post overshoot development.

**Fig. 7.**
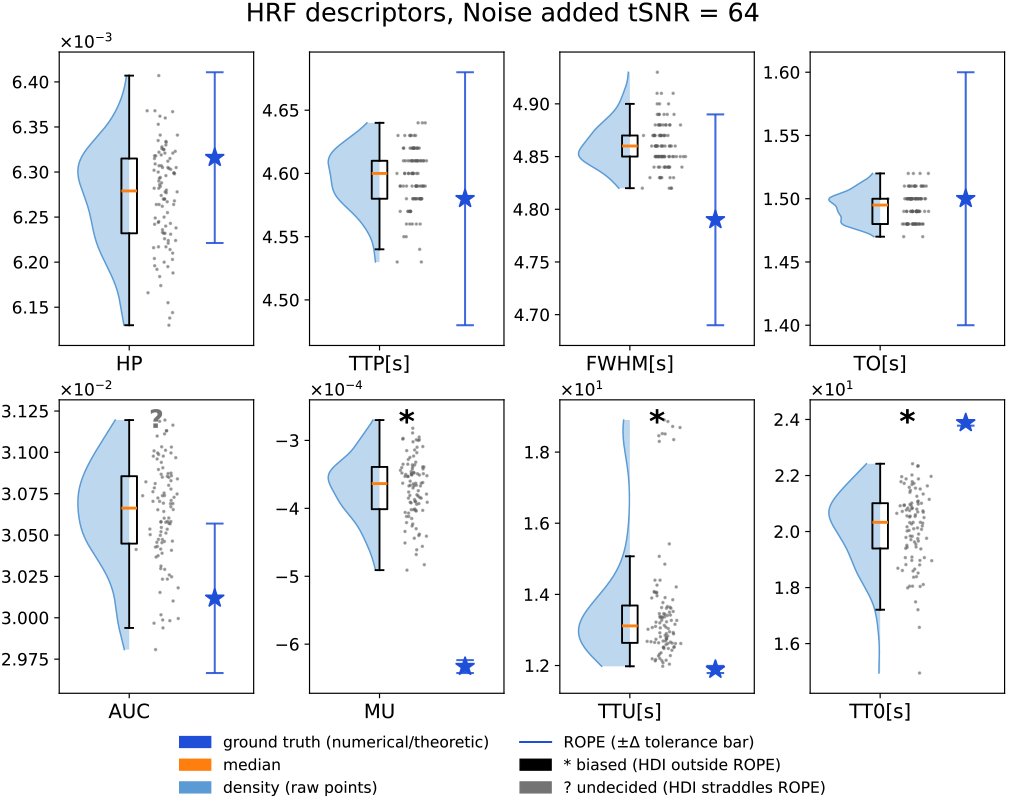
The noise added (tSNR=64) ensemble recovers four of eight HRF descriptors to within the pre-specified tolerance. Descriptor distributions over n=100 independent PINN fits of the same gaussian noise added simulated BOLD signal, each panel a raincloud (half-violin density, interquartile box, orange median, jittered raw runs). Blue star signing the ground truth from the numerical HRF; vertical blue bar, its region of practical equivalence (ROPE; *±*0.1 *s* for the temporal descriptors, *±*1.5% of groun-thruth for HP, MU, and AUC). Verdicts follow the HDI-versus-ROPE rule applied to the 95% HDI of the analytic Student-t posterior on the bias: unmarked, practically equivalent; black *\**, biased; grey ?, undecided. The biased descriptors are mainly related to a recovery phase: MU, TTU, and TT0. The undershoot is *≈*4.2 × 10^*−*5^ a.u higher, and sets almost 1.715s latter. TT0 sets *≈*4 s early. Why the AUC is labed as undecided whith an *≈−*1.52% bias. The MU offset is resolvable only because *n* = 100 restarts sharpen the posterior, not because it is large in absolute terms; the TTU and TT0 offsets are large on the scale of the response itself. Per-panel axis multipliers differ.

### In Vivo Data

Because the *in vivo* data offer no ground truth for their state variables, model adequacy is quantified through the fitness of the BOLD reconstructions against the measured signals (Table 3).

**Table 3.** Regression performance metrics against *in vivo* data, left(non-ischaemic, tSNR=76) and right (ischaemic, tSNR=66) hemisphere BOLD signal. *R*^2^ the coefficient of determination; *L*_2*RE*_ relative error and *S*_*ρ*_ Spearman Correlation coefficient.Values are ensemble means *±SD*(*×*10^*−*3^), *n* = 100 random-initialisation fits per hemisphere.

| | $R^2$ | $L_{2RE}$<br>(value ± SD($\times 10^{-3}$)) | $S_\rho$ |
| --- | --- | --- | --- |
| Left BOLD(t) | $0.415 \pm 3.260$ | $0.765 \pm 2.137$ | $0.656 \pm 2.957$ |
| Right BOLD(t) | $0.522 \pm 8.994$ | $0.691 \pm 6.555$ | $0.765 \pm 1.307$ |

The regression model showed moderate agreement with the in vivo BOLD signals in both hemispheres (Table 3). Model performance was consistently higher for the right hemisphere, with an increase in the coefficient of determination (*R*^2^: 0.522 vs. 0.415), a lower relative error (*L*_2*RE*_: 0.691 vs. 0.765), and a stronger monotonic association between predicted and measured signals (Spearman’s *ρ*: 0.765 vs. 0.656); considering the tSNR difference (right = 66, left = 76), signal quality cannot solely explain the observed differences, which may reflect biological or modelling factors. The latent state variables and the BOLD reconstruction for both hemispheres (Fig. 8) locate the differences chiefly in the deoxyhaemoglobin dynamics 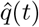. This interpretation must, however, be weighed against further studies.

**Fig. 8.**
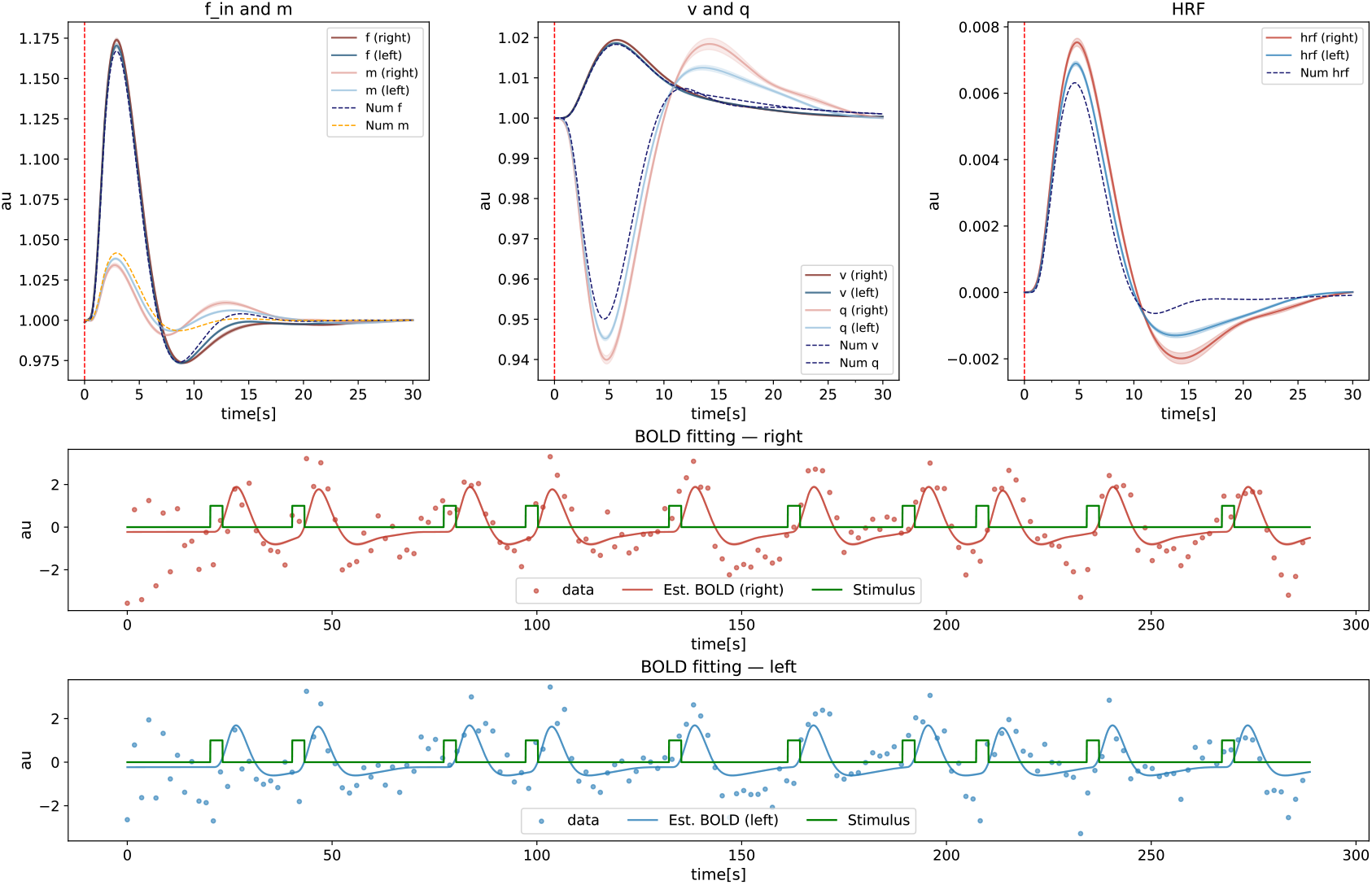
Hemispheric comparison of Balloon model state variables, HRF, and BOLD reconstruction, recovered from the in-vivo BOLD recordings. Two independent ensembles *n* = 100 PINN restarts each were trained, one per hemisphere; red denotes the right (ipsilesional, lower tSNR) and blue the left. Solid curves are across-restart means with a *±*1 SD band, whereas dashed lines show the numerical solution, shown for reference only, given that no ground truth exists for in-vivo data. First row, from left to right: *f* and *m* from Eq.3; the core of the Balloon Model venous volume *v* and deoxyhaemoglobin content *q* from Eq.2, the recovered HRF from Eq.1. The vertical red lines mark the beginning of the impulse needed to solve the Balloon equation system. Second and third rows, the ensemble-mean BOLD reconstruction (solid) against the measured time series (points) over the 300 s ten-block stimuli of 3*s* each (green), as shown in Eq.5. The right hemisphere shows the deeper deoxyhaemoglobin washout and the larger, later haemodynamic response (peak (7.54 *±* 0.12) *×* 10^*−*3^ versus (6.90 *±* 0.07) *×* 10^*−*3^ a.u., +9.3%; time to peak 4.788 *±* 0.034 versus 4.675 *±* 0.027 s; FWHM 5.267 *±* 0.050 versus 4.999 *±* 0.023 s;area under the positive section+14.7%; mean *±* SD, *n* = 100 per hemisphere). Both hemispheres overshoot the reference undershoot, the right by the larger margin (*−*1.99 *×* 10^*−*3^ versus *−*1.30 *×* 10^*−*3^ a.u.). The across-restart bands stay below 1.5% of each state’s dynamic range, so the hemispheric separation is far larger than the initialisation variability that could produce it. All quantities are in arbitrary units; time in seconds.

When studing the HRF estimation, the analysis is recast from a bias-against-truth assessment into a between-hemisphere comparison: the Bayesian estimation with a ROPE described in Sec. is applied to the difference 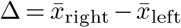 of each descriptor, where *n* = 100 random-initialisation fits are retained per hemisphere. The posterior of Δ, its 95% HDI, and the mass inside the descriptor-specific ROPE are reported in Table 7 (Appendix 4) and the per-descriptor distributions in Fig. 9.

**Fig. 9.**
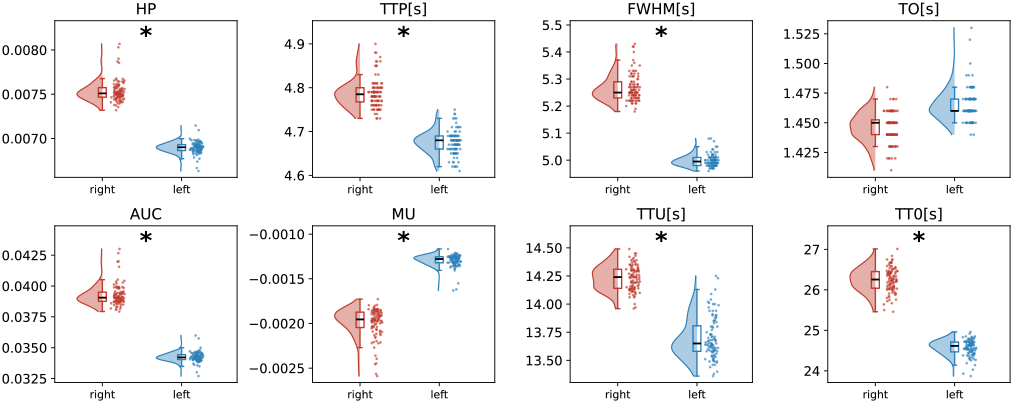
Per-descriptor distributions of the eight HRF descriptors for the right (ischaemic, red) and left (control, blue) hemispheres, each from *n* = 100 random-initialisation fits (raincloud: half-violin kernel-density estimate, narrow box, and jittered raw points). A black asterisk (*\**) marks descriptors whose between-hemisphere difference is different beyond practical equivalence under the Bayesian model; descriptors that are practically equivalent (only TO) carry no mark. The region of practical equivalence and the credible interval of the difference concern the between-group difference 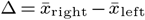 and are reported in Table 7(Appendix 4) rather than drawn on the group distributions.

Six of the eight descriptors differed between hemispheres beyond practical equivalence (HDI wholly outside the ROPE), whereas TTP and TO were practically equivalent (HDI wholly inside the *±*0.1 s ROPE), their interquartile ranges being the only ones to overlap in Fig. 9. The differences describe a broader and slower haemodynamic response on the right side: the HRF was wider (FWHM +0.165 s, reached its post-stimulus undershoot later (+0.484 s) and had a delayed TT0 (+1.332 s). Its amplitude was correspondingly greater, with a HP (+6.6%), a larger AUC (+9.8%) and a lower MU (Δ = *−*3.7 *×* 10^*−*4^). As in the simulated experiments, each descriptor is estimated independently from interval rather than accept/reject decisions, so no multiple-comparison correction is applied.

## Discussion

Estimation of the HRF from fMRI is a non-invasive method to investigate the brain’s haemodynamic response to task-driven neuronal activation. Phenomenological approaches (4, 14–16, 42) have been helpful so far; however, they lack a suitable biophysical correlate.

Physics-informed neural networks (PINNs), sometimes also referred to as theory-trained neural networks, incorporate mechanistic constraints (e.g., differential equations) directly into neural network training (35). Introduced by Raissi *et al*. (24), PINNs leverage the universal approximation capabilities of ANNs while penalising violations of the assumed governing equations, making them well suited for solving differential equations (forward problem) and performing parameter inference (backward problem) (22, 35, 43).

Our PINN approach to HRF estimation using block design fMRI data, can be cast as an optimisation problem that does not require prior knowledge of gamma probability distributions or suitable priors for Bayesian inference; instead, we used the Balloon model equations, constraining the regression to avoid overfitting to noisy data, while also providing a plausible *ad hoc* biophysical explanation from which to estimate haemodynamic responses using in vivo clinical data.

This approach relies on a balance between the weights assigned to physical knowledge and data. In our work, we show that for simulated signals with tSNR values similar to those of *in vivo* data acquired by our team, with a standard 1.5 T scanner under clinical conditions, we are able to retrieve the theoretical ground truth with a proportion (*ω*_*data*_*/ω*_*ode*_*/ω*_*ic*_*/ω*_*reg*_) of 0.60 : 0.40 : 1.0 : 1.0. When the same proportion is applied to clinical data, the retrieved state variables deviate from the vanilla solution (forward problem) of the balloon model equations, hinting at elements of a dynamic specific to the data.

Unlike our simulations with additive Gaussian noise, real fMRI data contains thermal and physiological noise, motion artefacts, and neural variability that simulated data do not; our results suggest that the PINN is not overfitting to noise. Despite a noisier target, the PINN achieves its best performance (goodness-of-fit) on right side, so signal quality alone does not account for the advantage.

The clinical applications of the present paradigm are not immediate; first of all, the training data came from a specific experimental design, and even when PINNs can be used to perform parameter estimation, inverse problems are known to admit multiple solutions; thus our results constitute one of many sets of state variables reconstructing the BOLD signal given our parametrisation, regularisation or variations of loss functions, its components or scalar weights determination might help with this endeavour.

Better signal quality would ordinarily be expected to ease the fit, so the higher *R*^2^ and Spearman correlation obtained on the noisier side ( right hemisphere tSNR = 66 vs. 76 on the left hemisphere)are the more notable for running counter to that expectation.

Our input is a scalar coordinate (normalised time) feeding a coordinate MLP that must represent four coupled state trajectories, spanning disparate timescales; structures are concentrated in the first third of the domain through early transients and slow recovery. Plain multilayer perceptrons are known to exhibit a bias toward low-frequency components (44, 45), motivating both the random weight factorisation of Wang et al. (37, 38) and our choice of a normalised activation. Unlike other smooth candidates, Softmax couples the units within each layer through a sum-to-one constraint, yielding a distribution over units rather than independent scalars. Post hoc, we find that the resulting representation expands substantially with training: the effective rank of the third-layer output, measured over the training grid, rises from *≈* 1.3 at initialisation to 3.79 after training. Since a linear map cannot increase the rank of its input, this expansion is attributable to the nonlinearity. We do not, however, separate the contributions of the factorisation and the activation, so the observed effect may operates through their training dynamics, and its disentanglement would require an ablation we did not perform. A systematic comparison against other smooth activations (tanh, sin, GELU) is likewise left to future work; the present study should be regarded as a proof of concept.

The results we present here were obtained under computational conditions that support iterative training and repeated model evaluation in a short time, resources that may not be routinely available in standard clinical environments.

## Conclusions

When constrained to solve the Balloon Windkessel equations, our PINN-based framework, in combination with fMRI data at a conventional magnetic field (1.5 T) and TR(1.75 s), estimates not only a haemodynamic response function, with descriptors in line with the literature, but also a plausible explanation, based on the model’s state variables. This estimation method produces physiologically consistent and personalised results, as it can be trained on single-subject data. To our knowledge, this is the first study to apply a single PINN to a highly nonlinear differential equation system, including the BOLD equation and convolution, in an indirect training. Beyond model refinement, future work could broaden its applicability by including a more extensive participant pool and diverse fMRI paradigms, while improving robustness through sensitivity analyses and modifications to the network architecture.

## Author Contributions

Rodrigo H. Avaria: Conceptualisation, Data Curation, Formal Analysis, Investigation, Methodology, Software, Validation, Visualization, Writing *−* Original Draft Preparation; David Ortiz: Methodology, Writing *−* Review & Editing; Javier Palma*−*Espinosa: Methodology, Visualization, Writing *−* reviewing and editing; Astrid Cancino: Data Curation, Writing *−* Review & Editing; Pablo Cox: Data Curation, Writing *−* Review & Editing; Rodrigo Salas: Conceptualisation, Funding Acquisition, Supervision, Writing *−* Review & Editing; Steren Chabert: Conceptualisation, Funding Acquisition, Project Administration, Resources, Supervision, Writing *−* Review & Editing.

## ACKNOWLEDGEMENTS

The authors used Claude Science IA to optimise the experiments’ code and to discuss and refine ideas regarding the data analysis (). Grammar, rhetoric, and pleonasm were assessed jointly with Claude Science and Grammarly to evaluate the readability of the finished document. All content was reviewed and edited by the authors, who take full responsibility for the final work.

The authors would like to acknowledge funding from the National Agency for Research and Development (ANID) through FONDECYT Grants No. 1231268 and No. 3250439, and the ANID–Millennium Science Initiative Programme ICN2021_004.

## Supplementary Note 1: Balloon Model Constants

The numerical values used for the Balloon–Windkessel constants and for the integration of Eq. (2) and Eq. (3) throughout this work are collected in Table 4; they follow (11, 20) unless stated otherwise.

**Table 4.** Model Variables and Typical Values according to (11, 20)

| Variable | Value |
| --- | --- |
| $\lambda_f$ | 0.2 |
| $\lambda_m$ | 0.05 |
| $\kappa$ | 1/1.54 |
| $\gamma$ | 1/2.46 |
| $dt$ (time step) | 0.01 |
| $\tau_{MTT}$ | 30 |
| $\tau$ | 20 |
| $\alpha$ | 0.4 |
| $V_0$ | 0.03 |
| $\vartheta_0$ | 40.3 |
| $E_0$ | 0.32 |
| $TE$ | 0.06 |
| $\epsilon$ | 1.43 |
| $r_0$ | 25 |

## Supplementary Note 2: Residuals and Time Standardisation

Before computational implementation, it is important to note that the orders of magnitude of the residuals from equations Eq. (3) and Eq. (2) (e.g. *f, m, v, q ~* 10^*−*2^) differ significantly from the difference between data and Eq. (5) (e.g. *Ŷ ~* 10^0^); and from the residuals against the initial conditions (e.g. *f* (*t < t*_*I*_); *m*(*t < t*_*I*_); *v*(*t < t*_*I*_), *q*(*t < t*_*I*_) ~ 10^*−*6^). During gradient descent optimisation, this disparity introduces complications, as it directly modulates the magnitude of gradients for the loss function *L*_*tot*_ during backpropagation.

Thus, we add a scalar weighting to level both *L*_*eq*_ and *L*_*ic*_ to the magnitude of *L*_*data*_ (36). Before training, we standardise the input features by mapping the HRF’s physical time domain [1, 30] to the [ *−*1.73, 1.73] interval with unit variance, a well-established technique for improving neural network convergence. Thus

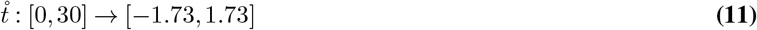

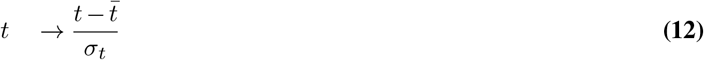

Where 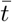, *σ*_*t*_ are the mean and standard deviation of the original temporal coordinate. Then the change of Variables Theorem compels us to rewrite the differential equations accordingly

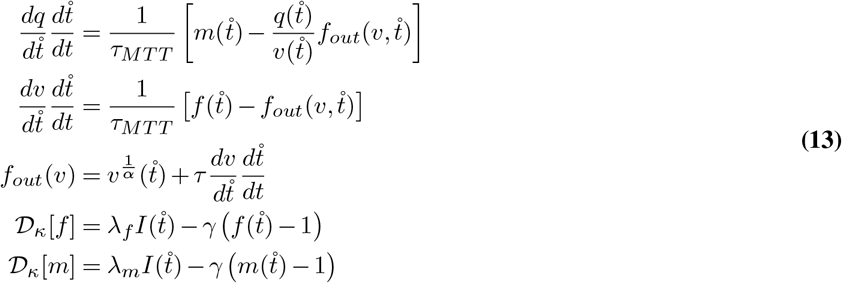

where the second-order operator is

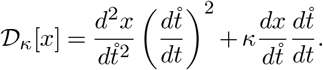

## Supplementary Note 3: Physics Regularisation

Previous results, not shown in this paper, showed greater standard deviation around the HRF, and a second family of functions minimising the total loss around 10% of training instances. Therefore we introduce a physics regularisation term. Starting from Eq. (2)

**Table 5.**
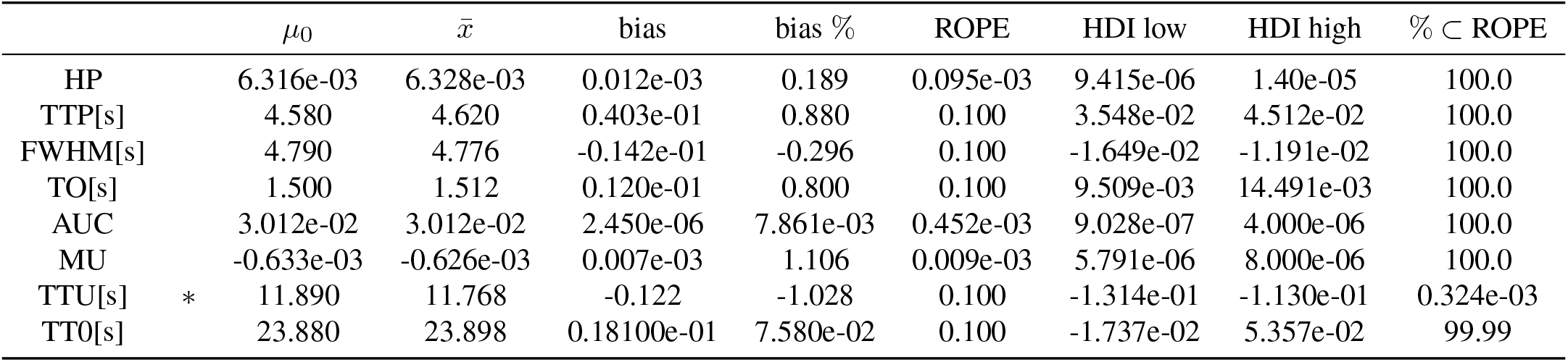
Bayesian practical-equivalence assessment of HRF descriptor recovery (noiseless ensemble, *n* = 100 random-initialisation fits). For each of the eight HRF descriptors, the bias 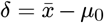 is the difference between the ensemble mean 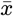 and the ground-truth value *µ*_0_ (obtained by applying the descriptor operator to the numerical HRF); *bias %* expresses this relative to *µ*_0_. The bias posterior is the closed-form Student-*t* distribution obtained under a Normal likelihood with the non-informative Jeffreys prior, 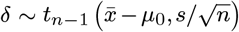; HDI_low_ and HDI_high_ bound 95% its highest-density interval. Δ is the Region of Practical Equivalence (ROPE) half-width, set to *±*0.1 s for the temporal descriptors (TTP, FWHM, TO, TTU, TT0) and *±*1.5% of |*µ*_0_| for the dimensionless/amplitude descriptors (HP, AUC, MU). % *⊂ ROPE* is the posterior mass within [*−*Δ, Δ]. The *\** marks the verdict following Kruschke’s HDI-and-ROPE rule: no sign if the HDI lies entirely inside the ROPE (therefore *equivalent*), *\** if entirely outside (*biased*), and *undecided* (?) if it straddles a limit. Seven descriptors are recovered to within acquisition resolution; time-to-undershoot (TTU) shows a small but well-resolved *−*0.122 s (*−*1.0%) early bias whose 95% HDI, [*−*0.131, *−*0.113] s, falls wholly below the *−*0.1 s limit (*<* 0.01% posterior mass in the ROPE). Because each descriptor is estimated independently and intervals rather than accept/reject decisions are reported, no multiple-comparison correction is applied; *n* indexes optimiser restarts, so the posterior describes the bias of the ensemble procedure.

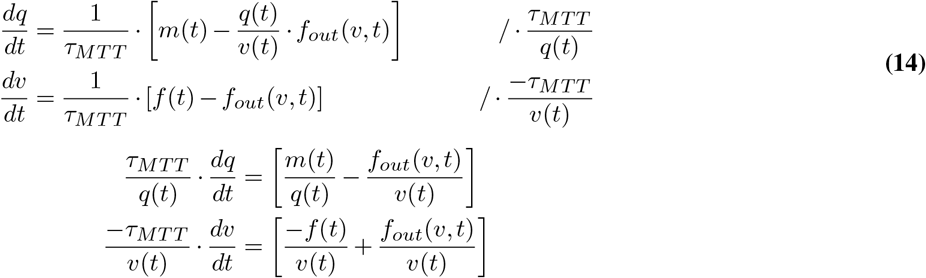

Now we can eliminate 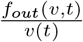 by adding the equations.

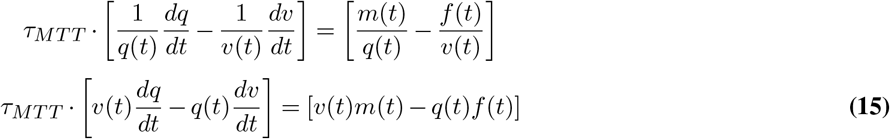

In Eq. (15) we get a direct expression relating all the outputs from the neural network, thus we use this equality as a regulariser.

## Supplementary Note 4: Tables

**Table 6.**
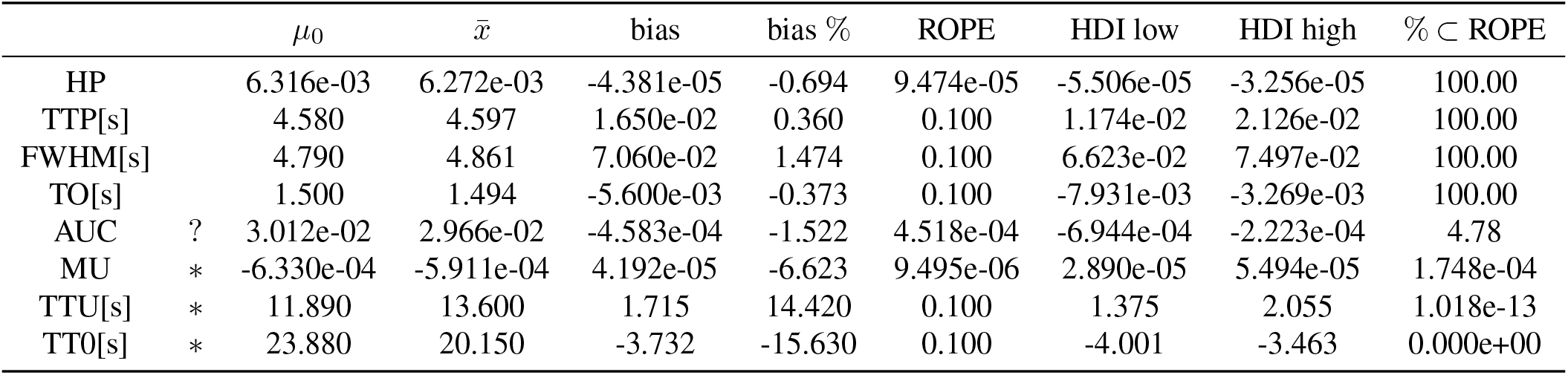
Bayesian practical-equivalence assessment of HRF descriptor recovery (noise-added ensemble, *n* = 100 random-initialisation fits). For each of the eight HRF descriptors, the bias 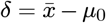 is the difference between the ensemble mean 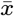 and the ground-truth value *µ*_0_ (obtained by applying the descriptor operator to the numerical HRF); *bias %* expresses this relative to *µ*_0_. The bias posterior is the closed-form Student-*t* distribution obtained under a Normal likelihood with the non-informative Jeffreys prior, 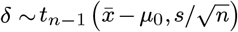; HDI_low_ and HDI_high_ bound 95%its highest-density interval. Δ is the Region of Practical Equivalence (ROPE) half-width, set to *±*0.1 s for the temporal descriptors (TTP, FWHM, TO, TTU, TT0) and *±*1.5% of |*µ*_0_| for the dimensionless/amplitude descriptors (HP, AUC, MU). % *⊂ ROPE* is the posterior mass within [*−*Δ, Δ]. The *\** marks the verdict following Kruschke’s HDI-and-ROPE rule: no sign if the HDI lies entirely inside the ROPE (therefore *equivalent*), *\** if entirely outside (*biased*), and *undecided* (?) if it straddles a limit. Under added measurement noise, four descriptors (HP, TTP, FWHM, TO) remain recovered to within acquisition resolution, whereas AUC becomes *undecided* (its 95% HDI, [ *−* 6.94, *−* 2.22] *×* 10^*−*4^, straddles the *±*4.58 *×* 10^*−*4^ limit; 2.8% posterior mass in the ROPE) and three descriptors are *biased* : time-to-undershoot (TTU) is late by +1.72 s (+14.4%, HDI [1.38, 2.06] s), time-to-zero (TT0) early by *−*3.73 s (*−*15.6%, HDI [*−*4.00, *−*3.46] s), and MU shows a small-magnitude but well-resolved shift (+4.19 *×* 10^*−*5^; large relative value reflecting the near-zero, signed ground truth). Because each descriptor is estimated independently and intervals rather than accept/reject decisions are reported, no multiple-comparison correction is applied; *n* indexes optimiser restarts, so the posterior describes the bias of the ensemble procedure.

**Table 7.** Bayesian assessment of hemispheric differences in HRF descriptors between the right-ischaemic and left-control ensembles (*n* = 100 random-initialisation fits per hemisphere; all runs retained). For each of the eight HRF descriptors the difference 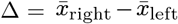 between the two hemispheric ensemble means is estimated under a Bayesian Behrens-Fisher model with unequal, never pooled, variances; Δ% expresses Δ relative to the mean of the two hemispheric means. HDI_low_ and HDI_high_ bound the 95% highest-density interval of the Δ posterior. Δ_ROPE_ is the Region of Practical Equivalence half-width, set to *±*0.1 s for the temporal descriptors (TTP, FWHM, TO, TTU, TT0) and *±*1.5% of the pooled-mean magnitude for the dimensionless/amplitude descriptors (HP, AUC, MU); % *⊂ ROPE* is the posterior mass within the Region. The *\** marks the verdict following Kruschke’s HDI-and-ROPE rule: no sign if the HDI lies entirely inside the ROPE (*equivalent*), *\** if entirely outside (*different*), ? and if it straddles a limit (*undecided*). Seven descriptors differ to a practically meaningful degree between hemispheres: the right-ischaemic HRF is broader (FWHM +0.268 s, +5.2%) with a later undershoot (TTU +0.536 s, +3.8%) and a later return to baseline (TT0 +1.677 s, +6.6%), and shows elevated amplitude (HP +8.8%, AUC +13.68%) rising slightly early (TTP +0.112 s, +2.4%); this TTP difference exceeds the ROPE by a small margin (HDI [0.104, 0.121] s) and should be read as a resolved but sub-acquisition-resolution shift rather than a physiologically large one. Only the onset descriptor TO (*−*0.019 s, *−*1.3%) is practically equivalent between hemispheres, its HDI lying wholly within the *±*0.1 s ROPE. The MU relative difference (*−*42.2%) is inflated by a near-zero, signed mean and should be read from the absolute Δ (*−*6.9 *×* 10^*−*4^) and its HDI rather than the percentage. Because interval estimates rather than accept/reject decisions are reported and each descriptor is assessed independently, no multiple-comparison correction is applied; *n* indexes optimiser restarts, so the posterior describes the difference between the two ensemble procedures.

| | | $\bar{x}_{\text{right}}$ | $\bar{x}_{\text{left}}$ | $\Delta$ | $\Delta \%$ | ROPE | HDI low | HDI high | $\% \subset \text{ROPE}$ |
| --- | --- | --- | --- | --- | --- | --- | --- | --- | --- |
| HP | * | 7.535e-03 | 6.897e-03 | 6.383e-04 | 8.8 | 1.082e-04 | 6.107e-04 | 6.658e-04 | 1.577e-78 |
| TTP[s] | * | 4.788 | 4.675 | 1.123e-01 | 2.4 | 0.100 | 1.037e-01 | 1.209e-01 | 2.581e-01 |
| FWHM[s] | * | 5.267 | 4.999 | 2.677e-01 | 5.2 | 0.100 | 0.257 | 0.279 | 4.838e-62 |
| TO[s] |  | 1.446 | 1.465 | -1.900e-02 | -1.3 | 0.100 | -2.272e-02 | -1.525e-02 | 100.00 |
| AUC | * | 3.927e-02 | 3.424e-02 | 5.029e-03 | 13.68 | 5.513e-04 | 4.837e-03 | 5.221e-03 | 8.212e-85 |
| MU | * | -1.989e-03 | -1.296e-03 | -6.932e-04 | -42.20 | 2.464e-05 | -7.287e-04 | -6.576e-04 | 0.000e+00 |
| TTU[s] | * | 14.23 | 13.70 | 0.536 | 3.8 | 0.100 | 0.493 | 0.578 | 3.415e-44 |
| TT0[s] | * | 26.25 | 24.57 | 1.677 | 6.6 | 0.100 | 1.606 | 1.748 | 1.443e-93 |

